# Atto 643 Carboxy Selectively Labels Astrocytes with Minimal Oligodendrocyte Cross-Reactivity

**DOI:** 10.64898/2026.03.09.710411

**Authors:** Xiaoqian Ge, Chun-Li Zhang, Zhenpeng Qin

**Affiliations:** Department of Biomedical Engineering, University of Texas Southwestern Medical Center, Dallas, TX 75390, USA; Department of Mechanical Engineering, The University of Texas at Dallas, Richardson, TX 75080, USA; Department of Molecular Biology, University of Texas Southwestern Medical Center, Dallas, TX 75390, USA; Hamon Center for Regenerative Science and Medicine, University of Texas Southwestern Medical Center, Dallas, TX 75390, USA; Peter O’Donnell Jr. Brain Institute, University of Texas Southwestern Medical Center, Dallas, TX 75390, USA; Department of Bioengineering, The University of Texas at Dallas, Richardson, TX 75080, USA; Center for Advanced Pain Studies, The University of Texas at Dallas, Richardson, TX 75080, USA; Harold C. Simmons Comprehensive Cancer Center, University of Texas at Southwestern Medical Center, Dallas, TX 75390, USA

## Abstract

Astrocytes are critical regulators of brain homeostasis and circuit function, and sulforhodamine 101 (SR101) are widely used for astrocyte labeling. However, SR101 also labels a significant portion of oligodendrocytes. Here, we identified Atto 643 carboxy, a far-red fluorescent dye, as a highly specific marker for astrocytes with minimal oligodendrocyte labeling. Furthermore, Atto 643 carboxy demonstrates minimal nonspecific uptake in myelin sheaths, and hippocampal pyramidal neuron as reported for SR101. We demonstrate robust imaging of astrocyte morphology and distribution in vivo and in acute brain slices. Our findings establish Atto 643 carboxy as a highly specific small-molecule probe for astrocyte-selective imaging, offering a powerful tool for dissecting astrocyte structure, dynamics, and function in intact brain tissue.

## Introduction

Astrocytes are the most abundant glial cell type in the central nervous system (CNS), playing key roles in regulating blood flow, synaptic function, neurovascular coupling, and behaviors through extensive interactions with neurons, vasculature, and other glial populations^1-4^. Throughout these processes, their elaborate, star-shaped morphology remains highly plastic, with structural remodeling often reflecting neural activity, injury, or disease, and serving as a sensitive indicator of altered brain states^5-7^. Furthermore, astrocytes exhibit remarkable diversity both across and within CNS regions^1, 8^. These studies have sparked a surge of interest in how astrocytes generate diverse physiological and pathological responses, a central question we probe through their complex, specialized morphology using selective labeling strategies^9, 10^.

Although genetic and viral tools enable astrocyte-specific labeling^11^, organic dyes such as SR101 and its analogs (e.g., sulforhodamine B) are chemical dyes used for astrocyte labelling^12-15^. Due to its ease of use, cost- and time effectiveness, SR101 have become indispensable in astroglia research. SR101 can be combined with neuronal markers, vascular tracers, or calcium indicators to investigate astrocyte–neuron communication, gliovascular coupling, and morphological dynamics in vivo or ex vivo brain tissue^13, 16-18^. Furthermore, SR101 can be applied to rat models where genetic manipulation is not readily available, providing a convenient and non-genetic approach for labeling astrocytes^13^. However, 20-40% of SR101 labeled cells are oligodendrocytes in cortical layers^12^. This is likely due to dye leakage through the gap junction between astrocytes and oligodendrocytes^12^. Additionally, prolonged incubation of SR101 dye can lead to dye infiltration into the myelin sheaths thus limiting its cellular specificity^12^. SR101 has also been shown to enter neurons through synaptic activity–dependent endocytotic uptake, followed by retrograde transport within presynaptic neurons^19^. Due to these limitations, there is an unmet need to identify reliable markers or probes to target astrocyte.

Here we report a far-red dye Atto 643 carboxy as a selective astrocyte marker. We assessed its cell-type specificity, regional labeling properties, and compatibility with in vivo two-photon imaging. Our results demonstrate that Atto 643 carboxy provides robust and selective labeling of astrocytes in multiple brain regions, with minimal uptake by neurons, microglia, NG2 glia, or oligodendrocytes. Furthermore, Atto 643 carboxy does not label myelin sheaths and retains high specificity across the cortical layers and the hippocampus. Finally, we show that its uptake is mediated by the organic anion-transporting polypeptide 1C1 (OATP1C1) transporter, which renders its astrocyte specificity. These findings establish Atto 643 carboxy as a powerful chemical tool for astrocyte imaging in both basic and translational neuroscience research.

## Results

Atto 643 is structurally related to the well-characterized rhodamine-based dye Atto 647N (**Fig. S1A**) but features modifications that confer significantly improved water solubility. To assess its cellular labeling in the brain, we injected functionalized form with the carboxy group (Atto 643 carboxy) into the caudate putamen (CPu; **Fig. 1A**), and, after 30 minutes, prepared acute brain slices for two-photon imaging. Atto 643 carboxy injections resulted in labelling cells that have multiple branched processes extending from the soma, suggestive of astrocytic identity (**Fig. 1B**). Many labeled cells displayed endfoot-like structures which envelops blood vessels, as confirmed by counterstaining the vascular lumen with intravenous injection of DyLight 488–lectin (**Fig. 1C**). Upon higher magnification, we observed Atto 643 carboxy^+^ cells exhibiting characteristic astrocytic morphology, including the soma, primary branches, fine branchlets, and leaflets (**Fig. 1D**).

**Fig. 1.**
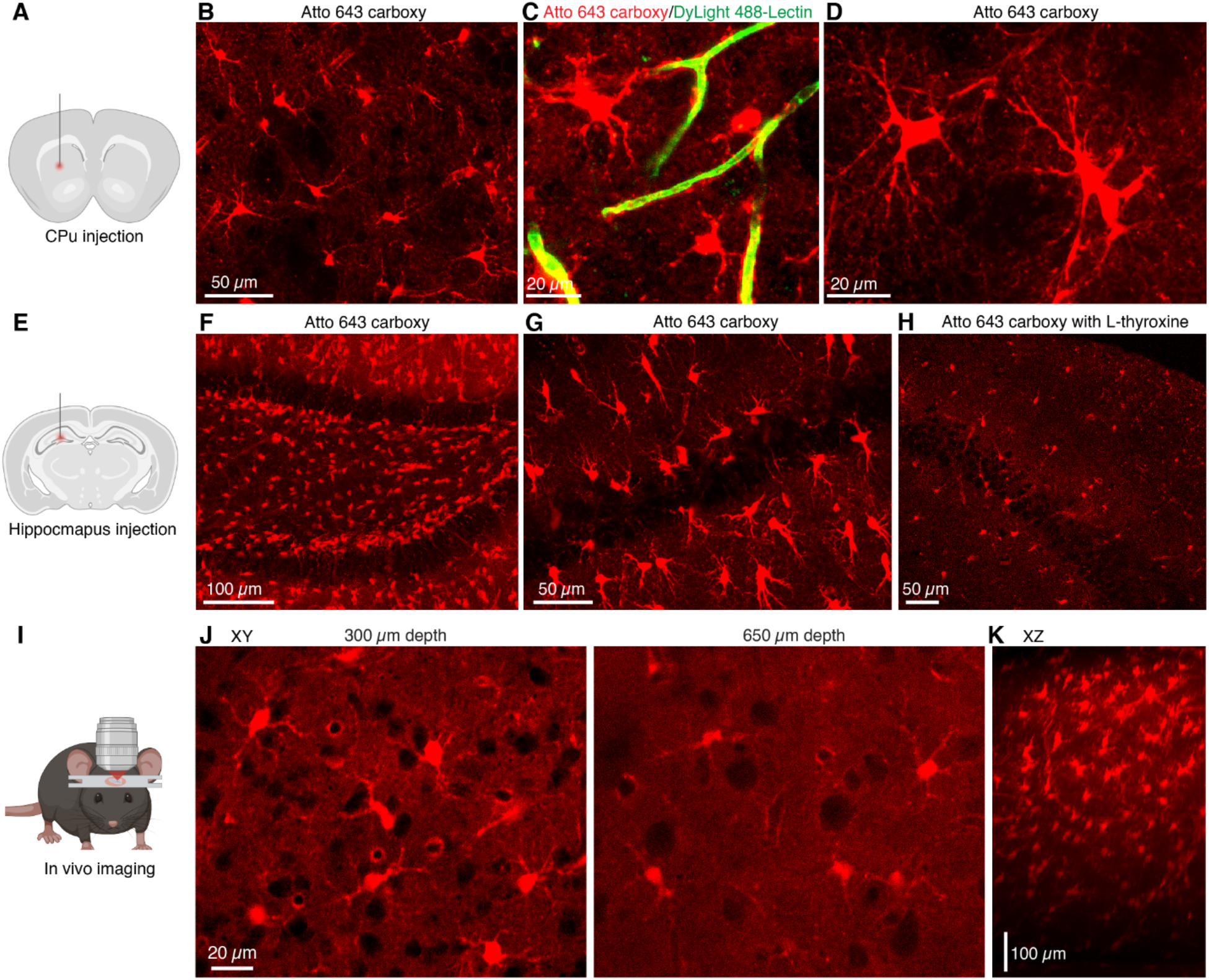
Atto 643 carboxy selectively labels astrocytes and supports intravital in vivo imaging. (**A**) Schematic of intracerebral injection of Atto 643 carboxy (1 µL, 50 µM) into the CPu, followed 30 min later by acute brain slice preparation for two-photon imaging. (**B**) Representative two-photon image showing Atto 643 carboxy^+^ cells in the CPu. (**C**) Intravenous injection of DyLight 488-lectin highlights blood vessels, confirming Atto 643 carboxy labeling of perivascular endfeet. (**D**) High-magnification two-photon image (maximum-intensity projection) reveals classical astrocytic features, including soma, primary branches, fine branchlets, and leaflets. (**E**) Illustration of hippocampal injection (0.4 µL, 50 µM) for two photon acute slice imaging. (**F**) Representative two-photon image (maximum-intensity projection) showing Atto 643 carboxy labeling in the dentate gyrus (DG) and (**G**) in CA1. (**H**) Co-injection of Atto 643 carboxy with L-thyroxine showing astrocyte labeling is markedly reduced in CA1. (**I**) Schematic of head-fixed in vivo two photon imaging setup following barrel cortex injection of Atto 643 carboxy. (**J**) Representative in vivo two-photon images showing Atto 643 carboxy labeling at depths of 300 µm and 650 µm. (**K**) Side-view projection (maximum-intensity, XZ) of Atto 643 carboxy-labeled cortical region, demonstrating up to 700 µm imaging depth.

Next, we assessed whether astrocyte labeling was unique to the carboxy-functionalized form of Atto 643 or could be replicated by other functional groups. To this end, we injected Atto 643 azide, Atto 643 biotin, and the structurally related Atto 647N carboxy into the CPu and imaged the tissue using two-photon microscopy. None of these variants labeled cells with characteristic astrocytic morphology (**Fig. S1B-E**), indicating that astrocyte labeling is specific to the carboxy-functionalized dye. We then examined regional specificity by injecting Atto 643 carboxy into the hippocampus, cortex, and brainstem. In the hippocampus and cortex, Atto 643 carboxy robustly labeled cells with astrocytic morphology (**Fig. 1E–F** and **Fig. S2A**), whereas no labeling was observed in the brainstem (**Fig. S2B**).

We tested inhibiting Atto 643 carboxy uptake in hippocampal astrocytes. Previous report suggests that SR101 uptake involves the thyroid hormone transporter OATP1C1^20^. To test this, L-thyroxine (T4), a known OATP1C1 inhibitor, was co-applied with Atto 643 carboxy into the hippocampus. Two-photon imaging performed 30 minutes post-application revealed a substantial reduction in both the number of labeled cells and fluorescence intensity compared to Atto 643 carboxy alone (**Fig. 1G–H**). The inhibited uptake in the hippocampus following OATP1C1 blockade, along with the absence of labeling in the brainstem, mirrors the pattern observed with SR101, suggesting that Atto 643 carboxy may act as a functional analog of sulforhodamine dyes.

Next, we checked if Atto 643 carboxy can be used in astrocyte labelling for live in vivo imaging. To this end, we injected the animals with Atto 643 carboxy in the barrel cortex and performed head-fixed two-photon imaging through a cranial window (**Fig. 1I**). We observed the labeling of a distinct subset of cortical cells, while large-soma, unlabeled cells were presumed to be neurons. Labeled cells displayed multiple processes and vessel-associated endfoot-like structures at all imaged depths (**Fig. 1J**), closely matching the astrocytic morphology seen in acute brain slices. The signal remained sharply resolvable to 700 µm below the pial surface (**Fig. 1K)**, indicating that Atto 643 carboxy enables deep, high-resolution in vivo imaging of astrocytes in cortical networks.

Our results thus far indicate the astrocyte labelling using Atto 643 carboxy in wild type mice. In order to verify the astrocyte specificity of Atto 643 carboxy and directly compare its labeling profile with SR101, we employed a transgenic mice model in which astrocytes express enhanced green fluorescent protein (EGFP) under the control of the aldehyde dehydrogenase 1 family member l1 (Aldh1l1) promoter (Tg(Aldh1l1-EGFP)). We injected either Atto 643 carboxy or SR101 into the CPu or cortex and assessed colocalization between dye-labeled cells (Atto 643^⁺^ or SR101^⁺^) and EGFP^⁺^ astrocytes in the acute brain slices using two photon imaging. Nearly all Atto 643 carboxy^⁺^ cells colocalized with EGFP^⁺^ cells, whereas a fraction of SR101^⁺^ cells lacked EGFP co-labeling (**Fig. 2A-B**). Quantitative analysis confirmed significantly higher colocalization for Atto 643 carboxy compared to SR101 in both the CPu (91.8 ± 7.1% vs. 35.7 ± 11.3%; p<0.001; n=6) and cortex (83.1 ± 11.5% vs. 52.0 ± 9.0%; p<0.001; n=6) (**Fig. 2C-D**). Furthermore, no Atto 643 carboxy labeling was observed in neurons or microglia (**Fig. 3A-B**), as confirmed by co-labeling with AAV9-hSyn-EGFP, which drives EGFP expression under the neuron-specific human synapsin (hSyn) promoter in wild-type mice, and with Tmem119-EGFP transgenic mice, which express EGFP specifically in microglia under the Tmem119 promoter. We assessed cross reactivity of Atto 643 carboxy dye in oligodendrocyte-lineage cells using NG2-DsRedBAC transgenic mice, which express DsRed in NG2 glia and vascular pericytes. Atto 643 carboxy showed minimal co-labeling with DsRed^⁺^ NG2 cells (**Fig. 3A-B**), compared to extensive NG2 co-labeling by SR101^12^. Together, these results demonstrate that Atto 643 carboxy is highly specific for astrocytes and labels them with substantially greater efficiency than the widely used SR101 dye.

**Fig. 2.**
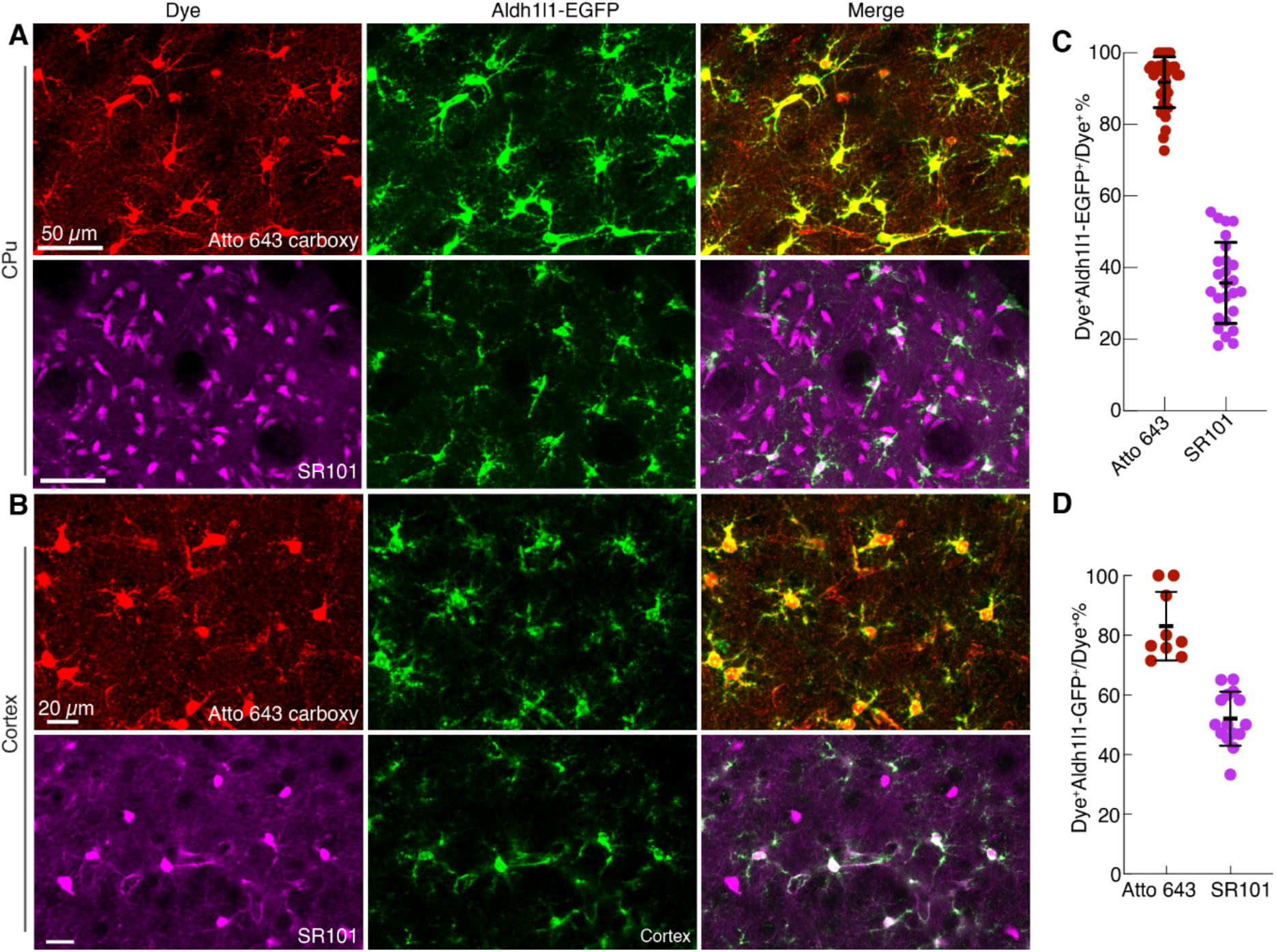
Astrocyte-specific labeling with Atto 643 carboxy. (**A** and **B**) Two-photon images of acute brain slices from Aldh1l1-EGFP transgenic mice demonstrate strong colocalization between Atto 643 carboxy^⁺^ cells and EGFP^⁺^ astrocytes in the CPu (**A**) and cortex (**B**), while SR101 shows partial overlap. Images are maximum-intensity projections and representative of 6 biological replicates. (**C** and **D**) Quantification of colocalization between dye^⁺^ cells and EGFP^⁺^ astrocytes in the CPu (**C**) and cortex (**D**), presented as the percentage of dye^⁺^ cells that are also EGFP^⁺^ (mean ± s.d.; each point represents one slice; n = 6 mice).

**Fig. 3.**
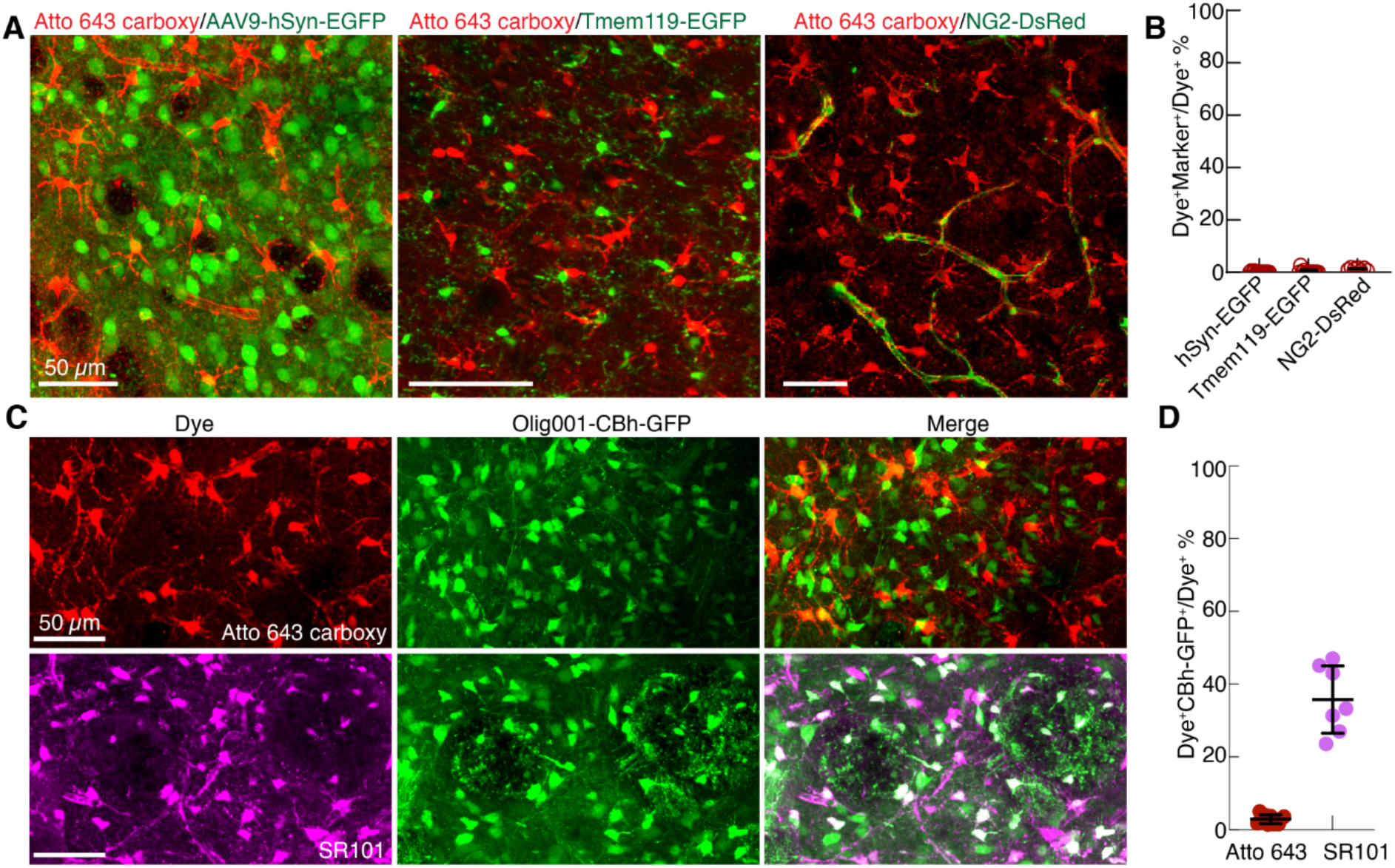
Minimal labeling of non-astrocytic cell types by Atto 643 carboxy. (**A**) Two-photon images showing Atto 643 carboxy labeling in brain slices from mice with neuronal (AAV9-hSyn-EGFP), microglial (Tmem119-EGFP), and oligodendrocyte-lineage (DsRed-NG2) labeling. (**B**) Quantification reveals that only 1.0 ± 0.7% of Atto 643 carboxy^⁺^ cells colocalize with EGFP^⁺^ neurons (6 Atto 643 carboxy^+^EGFP^+^ cells to 546 Atto 643 carboxy^+^ cell; n = 4 mice), 0.4 ± 0.2% of Atto 643 carboxy^⁺^ cells colocalize with EGFP^⁺^ microglia (2 Atto 643 carboxy^+^EGFP ^+^ cells to 1018 Atto 643 carboxy^+^ cells; n = 3 mice), and 1.3 ± 0.5% of Atto 643 carboxy^⁺^ cells colocalize with DsRed^⁺^ NG2 cells (14 Atto 643 carboxy^⁺^DsRed^+^ cells to 1106 Atto 643 carboxy^⁺^ cells; n = 3 mice). Each data point represents an individual acute slice. (**C**) Two-photon images of acute striatal brain slices from mice injected with Olig001-CBh-GFP virus, which labels oligodendrocytes, show minimal colocalization of Atto 643 carboxy^+^ cells with GFP^+^ oligodendrocytes, whereas SR101 labeling exhibits partial overlap. Images are presented as maximum intensity projections and are representative of 7 biological replicates. (**D**) Quantitative analysis of colocalization between dye^+^ cells with GFP^+^ oligodendrocytes. Data is presented as the percentage dye^+^ cells that colocalized GFP^+^ oligodendrocytes, relative to the total number of dye^+^ cells (mean ± s.d.; n = 7 Olig001-CBh-GFP injected mice).

One major drawback of the astrocytic marker SR101 is the “leakage” into oligodendrocytes^12^. To evaluate the Atto 643 carboxy’s oligodendrocyte cross reactivity and its relative performance to SR101, we utilized Olig001-CBh-GFP, a viral vector that selectively labels oligodendrocytes via GFP expression under the constitutive chicken β-actin (CBh) promoter^21^. Animals were first injected with the Olig001-CBh-GFP virus in the CPu, followed three weeks later by a second injection of either Atto 643 carboxy or SR101 into the same site. Thirty minutes post-injection, acute brain slices were prepared for two-photon imaging. Atto 643 carboxy exhibited minimal colocalization, with only 2.9 ± 1.2% of Atto 643 carboxy^+^ cells (24 Atto 643 carboxy^⁺^GFP^⁺^ cells to 798 Atto 643 carboxy^⁺^ cells, n =7 mice) overlapping with GFP^⁺^ oligodendrocytes, in contrast to SR101, which showed 35.8 ± 9.3% colocalization (296 SR101^⁺^GFP^⁺^ cells to 810 SR101^⁺^ cells, n =7 mice) (**Fig. 3C–D**). These findings indicate that Atto 643 carboxy has markedly lower cross-reactivity with oligodendrocyte cells compared to SR101 and highlights its specificity as an astrocytic marker.

In addition to oligodendrocyte cross reactivity, SR101 dye also suffers from nonspecific labeling in the CA1 region of the hippocampus. To evaluate whether Atto 643 carboxy shares this limitation, we injected the dye into the hippocampus, allowing diffusion across the CA1, CA3, and dentate gyrus (DG) subfields. Thirty minutes post-injection, acute slices were prepared to image dye labelling. Atto 643 carboxy selectively labeled cells with highly branched, star-like morphologies characteristic of protoplasmic astrocytes and did not label CA1 pyramidal neuron somata or dendrites (**Fig. 4A**). In contrast, SR101 prominently labeled CA1 pyramidal neurons, with only a minority of labeled cells displaying astrocytic morphology (**Fig. 4B**).

**Fig. 4.**
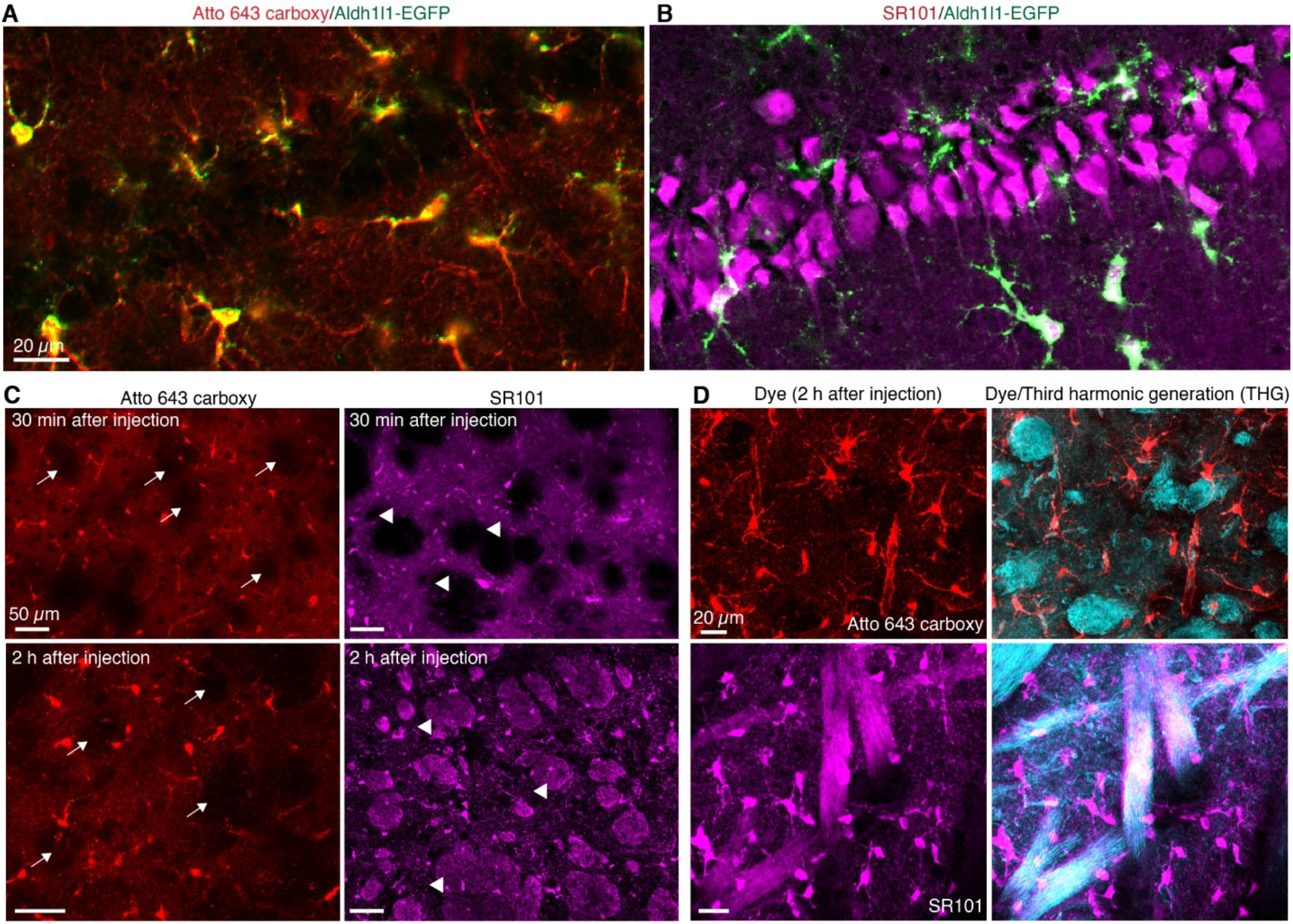
Atto 643 carboxy does not label hippocampal neurons and myelin sheath observed in SR101. (**A**-**B**) Two-photon images of the hippocampal CA1 region reveal robust SR101 labeling of pyramidal neurons, whereas Atto 643 carboxy shows no neuronal uptake. Images were repeated from 3 mice. (**C**) Two-photon imaging at 30 minutes post-injection revealed no myelin sheath labeling in the striosomes in the CPu with either Atto 643 carboxy or SR101. However, at 2 hours post-injection, SR101 exhibited prominent labeling of myelin sheaths, while Atto 643 carboxy remained absent from these structures. Striosomes are indicated with arrows in Atto 643 carboxy-labeled images and with arrowheads in SR101-labeled images. (**D**) Simultaneous two-photon fluorescence and third harmonic generation imaging indicate SR101 labeling of myelin sheaths at 2 hours post-injection, which is absent with Atto 643 carboxy. Images are shown as maximum intensity projections. Images were repeated from 4 mice.

To compare myelin sheath labeling by Atto 643 carboxy and SR101, we injected each dye into the CPu and examined acute brain slices at 30 min and 2 h post-injection, focusing on striosome regions where myelin is abundant. At 30 min, neither dye labeled myelin sheaths. By 2 h, SR101 displayed clear myelin sheath labeling, whereas Atto 643 carboxy remained absent from these structures (**Fig. 4C**). To confirm these observations, we conducted simultaneous two-photon fluorescence and third harmonic generation (THG) imaging, a dye-free method for visualizing myelin^22^. THG imaging revealed strong myelin signals in striosomes of the CPu that overlapped with SR101, but not with Atto 643 carboxy (**Fig. 4D**), confirming the lack of myelin labeling by Atto 643 carboxy.

## Discussion

In this study, we identified Atto 643 carboxy as a highly selective, cost and time efficient fluorescent marker for astrocytes. Atto 643 carboxy enables rapid astrocyte labeling following low-volume stereotaxic injection and supports high-resolution imaging of astrocyte morphology and spatial domains. Its compatibility with two-photon microscopy and far-red emission spectrum facilitates multiplexing with other fluorescent reporters, such as neuronal or vascular markers and Ca^2⁺^ indicators, thus expanding its utility in studies of tripartite synapses, gliovascular coupling, and astrocyte-neuron signaling in live preparations. Our data show superior results compared to the previously reported astrocytic dye SR101. While SR101 is widely used in astrocyte research for its simplicity and compatibility with live imaging, it suffers from substantial off-target labeling, particularly in oligodendrocytes and myelin sheaths, and nonspecific labels neurons in hippocampus. In contrast, Atto 643 carboxy demonstrates robust and specific labeling of astrocytes across multiple brain regions, with minimal labeling of neurons, microglia, NG2 glia, or oligodendrocytes.

Mechanistically, our findings suggest that Atto 643 carboxy shares a common uptake pathway with SR101, potentially involving the thyroid hormone transporter OATP1C1. The reduction in dye uptake following co-administration with L-thyroxine, an OATP1C1 inhibitor, supports this hypothesis. The structure of Atto 643 carboxy is unknown, however, the spectral and uptake similarities between Atto 643 carboxy and SR101 raise the possibility that the former is a sulforhodamine analog with improved physicochemical properties. Atto 643 carboxy’s increased molecular weight and reduced diffusion properties may underlie its enhanced specificity. One limitation of Atto 643 carboxy dye is its incompatibility with standard fixation protocols, which restricts its use in immunohistochemical assays. Like SR101, Atto 643 carboxy may require repeated administration for extended imaging sessions. Future studies aimed at optimizing its formulation or conjugating the dye to fixable carriers may further expand its versatility.

Collectively, our findings demonstrate a simple but powerful tool for visualizing astroglia, combining rapid and selective labeling with minimal off-target staining. This has enormous potential within the neuroscience research for studying astrocyte diversity, neurovascular interactions, monitoring dynamic responses, and astrocyte–neuron communication. Its specificity, compatibility with in vivo imaging, and potential for integration with multi-channel fluorescent approaches make it well-suited for dissecting the complex roles of astrocytes in brain physiology and disease.

## METHODS

### Animals

Wild-type C57BL/6 mice were purchased from Charles River Laboratory. Transgenic mice Aldh1l1-EGFP/Rpl10a JD130 (Strain: 030247), Tmem119-2A-EGFP (Strain: 031823), and NG2DsRedBAC (Strain: 008241) were purchased from Jackson Laboratory. All animals were used with age around 8-12 weeks with both sexes. Animal protocols adhered to the National Institutes of Health (NIH) guidelines and received approval from the institutional animal care and use committee at University of Texas Southwestern Medical Center.

### Reagents

Atto 643 carboxy, Atto 643 azide, Atto 643 biotin, and Atto 647N carboxy were purchased from ATTO-TEC GmbH (Siegen, Germany). SR101 (Catalog No. S359) and Dylight 488 lectin (Catalog No. L32470) were purchased from ThermoFisher Scientific, and L-thyroxine was purchased from Sigma-Aldrich (Catalog No. T2376). Atto 643 and SR101 dyes were dissolved in 1X PBS at a concentration of 10 mM, while Atto 647N dye and L-thyroxine were dissolved in dimethyl sulfoxide (DMSO) at the concentration of 5 mM. All the reagents were aliquoted and stored at -20 °C before use.

### Surgeries

Mice were anesthetized with isoflurane (5% for induction and 1.5% for maintenance) and secured in a stereotaxic apparatus using ear bars (Kopf Instruments). The body temperature was maintained at 34 °C using a thermostatically controlled heating pad and monitored with a rectal probe. Buprenorphine (1 mg/kg, subcutaneous) was administered for analgesia. Intracerebral injections were carried out using a Nanoliter 2020 injector (World Precision Instruments) fitted with a 30 μm diameter glass pipette prepared using a micropipette puller (Sutter Instrument Co., P-97).

The scalp was incised and retracted, and a cranial window was created using a dental drill. Intracerebral administration of Atto 643 dye or SR101 (50 μM, diluted in 1XPBS) into the cranial window was performed at specified stereotaxic coordinates corresponding to multiple brain regions including the cortex, caudate-putamen, hippocampus, and brainstem. Coordinates (from bregma point) for these brain regions were mediolateral (M/L, 2.0 mm), anteroposterior (A/P, -2.0 mm), and dorsoventral (D/V, 0.8 mm) for cortex; M/L 2 mm, A/P 0 mm, and D/V 3 mm for caudate-putamen; M/L 1.2 mm, A/P -1.5, and D/V 1.8 mm for hippocampus; and M/L 1 mm, A/P -6.5 mm, and D/V 4.5 mm for brainstem, respectively. 0.4 μL dye (0.3 nL/s) was injected into the cortex or hippocampus, and 1 μL dye (1 nL/s) was injected into caudate-putamen or brainstem. After infusions, the glass pipette was left in place for an additional 10 minutes to enable diffusions. Following the injection, the glass pipette was gently removed, and mice were sacrificed or housed before acute brain slice preparation.

To minimize the risk of edema and membrane disruption that could interfere with cell-type specificity analysis, intracerebral injections were performed using a total volume of less than 1 µL per mouse. The injection was delivered at a controlled rate not exceeding 1 nL/s to ensure minimal tissue disturbance.

### Adeno-associated virus (AAV) vector administration

To label neurons, AAV9-hSyn-EGFP (Addgene, 50465-AAV9; titer: 1.3 × 10^13^ vg/mL) was injected into the caudate-putamen at a dose of 1.3 × 10^9^ vg per mouse. Mice were housed for three weeks to allow for EGFP expression in neurons. To label oligodendrocytes, Olig001-CBh-GFP viral particles (titer: 1.1 × 10^13^ vg/mL), produced by the UT Southwestern Viral Vector Core, were injected into the caudate-putamen at a dose of 1.1 × 10^9^ vg per mouse. Mice were similarly housed for three weeks to ensure sufficient GFP expression in oligodendrocytes. Following the expression period, 1 μL of either Atto 643 carboxy or SR101 (50 μM in 1X PBS) was injected into the previously transduced region. After 30 minutes, animals were euthanized for acute brain slice preparation.

### Acute brain slice preparation

Acute coronal brain slices (300 μm thick) were prepared on a Leica VT1200 vibratome. Specifically, slicing was performed in ice-cold, modified artificial cerebrospinal fluid (ACSF) with reduced Na^+^ and Ca^2+^ and elevated Mg^2+^ concentrations (slicing ACSF: 72 mM NaCl, 2.5 mM KCl, 26 mM NaHCO_3_, 1.25 mM NaH_2_PO_4_, 0.5 mM CaCl_2_, 2 mM MgSO_4_, 24 mM glucose, and 75 mM sucrose; osmolarity ∼300 mOsm/L). Immediately after sectioning, slices were transferred to a recovery ACSF solution (124 mM NaCl, 5 mM KCl, 26 mM NaHCO_3_, 1.25 mM NaH_2_PO_4_, 1.5 mM CaCl_2_, 1.3 mM MgCl_2_, and 10 mM glucose; osmolarity ∼300 mOsm/L) and maintained in a brain slice keeper (Automate Scientific, BSK4). Slices were incubated at 34 °C for 30 minutes, followed by an additional 1-hour incubation at room temperature. Both slicing and recovery ACSF solutions were continuously bubbled with carbogen gas (95% O_2_/5% CO_2_).

### Acute brain slice imaging

Acute brain slices were transferred to an open-bath recording chamber mounted on a #1.5 coverslip (Fisher Scientific) and positioned within a microscope stage adapter (Warner Instruments, SA-20UU-AL). The recording chamber was integrated with a perfusion system consisting of a heating platform (Warner Instruments, PH-1), a peristaltic pump (Sigma-Aldrich, Z678406), and a vacuum system (Warner Instruments, 64-1940), enabling continuous ACSF perfusion to maintain slice viability during imaging. The heating platform was regulated by a dual-channel temperature controller (Warner Instruments, TC-344C), maintaining the chamber temperature at 34 °C. Oxygenated recovery ACSF was delivered through the peristaltic pump and further heated using an inline solution heater (Warner Instruments, SH-27B). The perfusion rate was maintained at 2 mL/min.

### Two photon microscopy

Two photon Imaging was performed using a commercial Olympus MPE-RS twin multiphoton microscope equipped with an Insight DS+ OL pulsed infrared (IR) laser (Spectra-Physics), providing tunable excitation wavelengths ranging from 680 to 1300 nm with a pulse width of 120 fs. Emitted signals were collected through water-immersion objectives of varying magnification: 10× (XLPLN10XSVMP, NA 0.6), 25× (XLPLN25XWMP2, NA 1.05), and 40× (LUMPLFLN40XW(F), NA 0.80). Emission light passed through a dichroic mirror (Olympus, DM760) and a short-pass filter (Olympus, FV30-SDM570), directing signals above 570 nm to GaAsP photomultiplier tubes (PMTs) and those below 570 nm to multi-alkali PMTs. Fluorophores were excited at their respective optimal wavelengths: Atto 643 at 1200 nm, DsRed and SR101 at 1000 nm, and GFP and DyLight 488-lectin at 920 nm. Emissions from Atto 643, DsRed, and SR101 were collected using the FV30-FOCY5 filter cube, with spectral separation at 650 nm into two channels: BA660–750 nm and BA575–645 nm. Emissions from GFP and DyLight 488-lectin were collected using the FV30-FGR filter cube with a detection range of BA495–540 nm. Laser power, electron-multiplying (EM) gain, and offset settings were optimized for each experiment to maximize fluorescence signal-to-noise ratio while minimizing spectral crosstalk. Imaging Z-stacks were acquired with a step size of 1 μm, covering depths of 50–200 μm in the acute brain slice. All images were acquired at a resolution of 1024 × 1024 pixels with a pixel size of 0.5 μm and processed using Olympus cellSens dimension software.

### Third harmonic generation imaging

Simultaneous two-photon fluorescence and third harmonic generation imaging (THG) were performed using an Olympus MPE-RS twin multiphoton microscope. Two-photon fluorescence imaging of Atto 643 carboxy or SR101-labeled cells in acute brain slice was conducted as described in the Two-photon microscopy section. For THG imaging, the excitation wavelength was set to 1250 nm, and emission was collected using the FV30-FVG filter cube. The emission signal was spectrally split at 485 nm into two detection channels: BA410–460 nm and BA495– 540 nm. The BA410–460 nm channel was used to capture the THG signal. Image processing was performed using the same workflow described for two-photon fluorescence imaging.

### In vivo two-photon imaging

To label astrocytes in the barrel cortex, 400 nL of Atto 643 carboxy (50 µM) was injected intracerebrally using stereotaxic coordinates: M/L 2.0 mm, A/P ™2.0 mm, and D/V 0.8 mm. Following the injection, a 3–4 mm circular cranial window centered on the injection site was outlined. The skull within the marked region was carefully removed using a microdrill, and the exposed surface was cleaned and dried. A 3 mm round coverslip was then placed over the craniotomy and sealed with biological adhesive, ensuring full coverage of the exposed area and minimizing bleeding throughout the procedure. A custom-designed steel head bar was affixed to the skull, and a circular moat was constructed around the cranial window using dental cement. Mice were placed on the imaging stage of the two-photon microscope and maintained under anesthesia with a portable isoflurane system. Body temperature was kept constant at 34 °C using a thermostatically controlled heating pad. During imaging, the moat was filled with pre-warmed ACSF, and imaging was performed using a 25× water-immersion objective (XLPLN25XWMP2, NA 1.05). Initial focusing was guided by visualization of unlabeled surface vessels. Atto 643 carboxy fluorescence was excited at 1200 nm, and emission was collected in the 660–750 nm range using the FV30-FOCY5 filter cube. Image stacks were acquired from the pial surface to deeper cortical layers and analyzed using Fiji software (National Institutes of Health, NIH). XY panels were generated from selected Z-depths, and XZ views were reconstructed via reslicing and side-view projection. No additional images processes were applied.

### Dye uptake inhibition

To investigate the inhibition of Atto 643 carboxy uptake, we adapted a previously reported protocol with minor modifications^20^. L-thyroxine was prepared at a concentration of 50 µM in 1× PBS, and 1 µL was injected into the hippocampus using the following stereotaxic coordinates: M/L 1.2 mm, A/P ™1.5 mm, and D/V 1.8 mm. One hour following L-thyroxine administration, 400 nL of Atto 643 carboxy (50 µM) combined with L-thyroxine (50 µM) was injected into the same site. Mice were then euthanized, and acute brain slices were prepared for two-photon imaging. For the control group, 1 µL of 1× PBS was injected into the hippocampus, followed by 400 nL of Atto 643 carboxy (50 µM) at the same coordinates and time interval.

### Image analysis

Image processing was initially performed using Fiji to convert raw data into TIFF format. Cell segmentation was carried out using Cellpose^23^, and regions of interest (ROIs) were extracted for each identified cell. These ROIs were subsequently imported back into Fiji for manual inspection and correction of any false segmentations. The number of dye^⁺^ and marker^⁺^ cells in each fluorescence channel was quantified using Fiji’s “analyze/measure” function. Colocalization between dye^⁺^ and marker^⁺^ cells was determined by identifying overlapping ROIs between the two channels. The number of double-positive cells (dye^⁺^ marker^⁺^) was counted and used to calculate labeling efficiency, defined as the ratio of dye^⁺^ marker^⁺^ cells to the total number of dye^⁺^ cells.

### Statistical analysis

All statistical analyses were performed using GraphPad Prism. Sample sizes (n) and corresponding p-values are reported in the figures, figure legends, or main text as appropriate. Data are presented as mean ± standard deviation (s.d.). Statistical significance was assessed using appropriate parametric or non-parametric tests, with a predefined significance threshold of α = 0.05.

## Acknowledgements

This study was sponsored by Cancer Prevention and Research Instituteof Texas (CPRIT) grants RP190278 and RP210236, Department of Defense grant W81XWH-21-1-0219 and American Heart Association grant 19CSLOI34770004, National Science Foundation (NSF) under grant number 2123971, National Institutes of Health (NIH) under grant numbers RF1NS110499 and R01MH140482, and Eugene McDermott Professorship at the University of Texas at Dallas. Schematic images were obtained from Biorender.com. We thank Dr. Steven Gray from UT Southwestern Viral Vector Core for preparing Olig001-CBh-GFP viral particles.

## Author Contributions

X.G. and Z.Q. generated the idea, and X.G. performed experiments and analyzed data. C.Z. helped with data analysis and design experiments. X.G. drafted the manuscript. Z.Q. and X.G. revised the manuscript. All authors contributed to the writing of this paper

## Notes

### Competing Interest Statement

The authors have declared no competing interest.

## Reference

1. Endo, F. et al. Molecular basis of astrocyte diversity and morphology across the CNS in health and disease. Science 378, eadc9020 (2022).

2. Hirrlinger, J. & Nimmerjahn, A. A perspective on astrocyte regulation of neural circuit function and animal behavior. Glia 70, 1554–1580 (2022).

3. Khakh, B.S. & Sofroniew, M.V. Diversity of astrocyte functions and phenotypes in neural circuits. Nat. Neurosci. 18, 942–952 (2015).

4. Verkhratsky, A., Nedergaard, M. & Hertz, L. Why are Astrocytes Important? Neurochem. Res. 40, 389–401 (2015).

5. Díaz-Castro, B., Robel, S. & Mishra, A. Astrocyte Endfeet in Brain Function and Pathology: Open Questions. Annu. Rev. Neurosci. 46, 101–121 (2023).

6. Lee, H.-G., Wheeler, M.A. & Quintana, F.J. Function and therapeutic value of astrocytes in neurological diseases. Nat. Rev. Drug Discov. 21, 339–358 (2022).

7. Acosta, C., Anderson, H.D. & Anderson, C.M. Astrocyte dysfunction in Alzheimer disease. J. Neurosci. Res. 95, 2430–2447 (2017).

8. Westergard, T. & Rothstein, J.D. Astrocyte Diversity: Current Insights and Future Directions. Neurochem. Res. 45, 1298–1305 (2020).

9. Kriegstein, A. & Alvarez-Buylla, A. The Glial Nature of Embryonic and Adult Neural Stem Cells. Annu. Rev. Neurosci. 32, 149–184 (2009).

10. Baldwin, K.T., Murai, K.K. & Khakh, B.S. Astrocyte morphology. Trends Cell Biol. 34, 547–565 (2024).

11. Hülsmann, S., Hagos, L., Heuer, H. & Schnell, C. Limitations of Sulforhodamine 101 for Brain Imaging. Front. cell. neurosci. 11 (2017).

12. Hill, R.A. & Grutzendler, J. In vivo imaging of oligodendrocytes with sulforhodamine 101. Nat. Methods 11, 1081–1082 (2014).

13. Nimmerjahn, A., Kirchhoff, F., Kerr, J.N.D. & Helmchen, F. Sulforhodamine 101 as a specific marker of astroglia in the neocortex in vivo. Nat. Methods 1, 31–37 (2004).

14. Appaix, F. et al. Specific In Vivo Staining of Astrocytes in the Whole Brain after Intravenous Injection of Sulforhodamine Dyes. PLoS One 7, e35169 (2012).

15. Schnell, C., Hagos, Y. & Hülsmann, S. Active Sulforhodamine 101 Uptake into Hippocampal Astrocytes. PLoS One 7, e49398 (2012).

16. Damisah, E.C., Hill, R.A., Tong, L., Murray, K.N. & Grutzendler, J. A fluoro-Nissl dye identifies pericytes as distinct vascular mural cells during in vivo brain imaging. Nat. Neurosci. 20, 1023–1032 (2017).

17. Watanabe, A., Guo, C. & Sjöström, P.J. The developmental profile of visual cortex astrocytes. iScience 26, 106828 (2023).

18. Tischbirek, C., Birkner, A., Jia, H., Sakmann, B. & Konnerth, A. Deep two-photon brain imaging with a red-shifted fluorometric Ca2+ indicator. Proc. Natl. Acad. Sci. U.S.A. 112, 11377–11382 (2015).

19. Cina, C. & Hochman, S. Diffuse distribution of sulforhodamine-labeled neurons during serotonin-evoked locomotion in the neonatal rat thoracolumbar spinal cord. J. Comp. Neurol. 423, 590–602 (2000).

20. Schnell, C. et al. The multispecific thyroid hormone transporter OATP1C1 mediates cell-specific sulforhodamine 101-labeling of hippocampal astrocytes. Brain Structure and Function 220, 193–203 (2015).

21. Powell, S.K. et al. Characterization of a novel adeno-associated viral vector with preferential oligodendrocyte tropism. Gene Ther. 23, 807–814 (2016).

22. Farrar, Matthew J., Wise, Frank W., Fetcho, Joseph R. & Schaffer, Chris B. In Vivo Imaging of Myelin in the Vertebrate Central Nervous System Using Third Harmonic Generation Microscopy. Biophys. J. 100, 1362–1371 (2011).

23. Stringer, C., Wang, T., Michaelos, M. & Pachitariu, M. Cellpose: a generalist algorithm for cellular segmentation. Nat. Methods 18, 100–106 (2021).

