## Supporting information for "Atto 643 Carboxy Selectively Labels Astrocytes with Minimal Oligodendrocyte Cross-Reactivity"

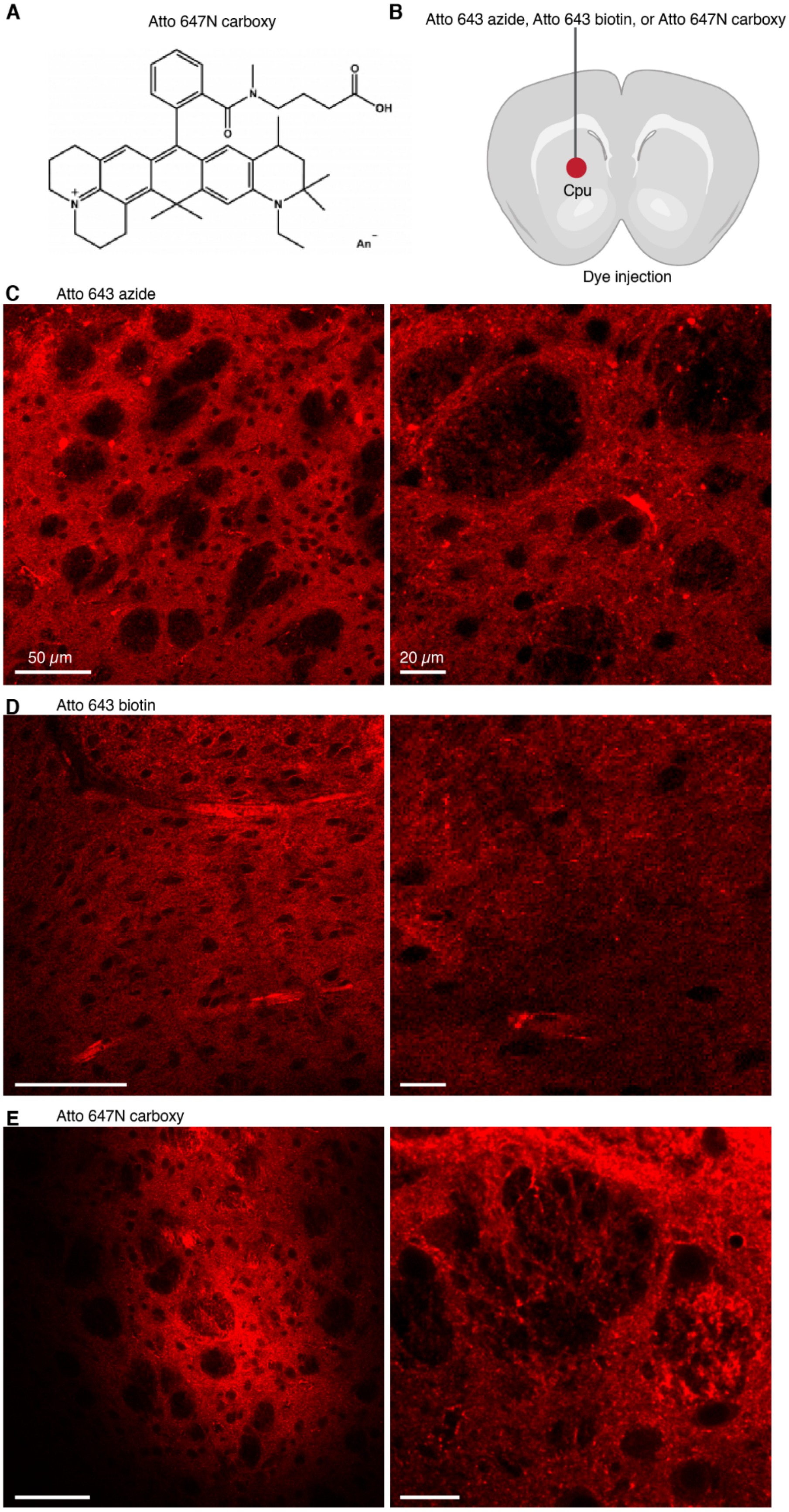
A
Atto 647N carboxy
B
Atto 643 azide, Atto 643 biotin, or Atto 647N carboxy
Cpu
Dye injection
C
Atto 643 azide
50 μm
20 μm
D
Atto 643 biotin
E
Atto 647N carboxy

**Fig. S1. Alternative Atto 643 variants and Atto 647N carboxy analog do not label astrocytes.** (**A**) Chemical structure of Atto 647N carboxy, a rhodamine-based dye related to Atto 643; structure and information obtained from the ATTO-TECH GmbH website. (**B**) Wild-type mice were injected into the caudate putamen (CPu) with 1 μL of 50 μM Atto 643 azide, Atto 643 biotin, or Atto 647N carboxy. Acute brain slices were prepared for two-photon imaging thirty minutes post-injection. (**C–E**) Two-photon imaging showed that none of the tested dyes significantly labeled astrocytes. Fluorescence signals were primarily extracellular or associated with blood vessels, with only minimal cellular uptake observed for Atto 643 azide.

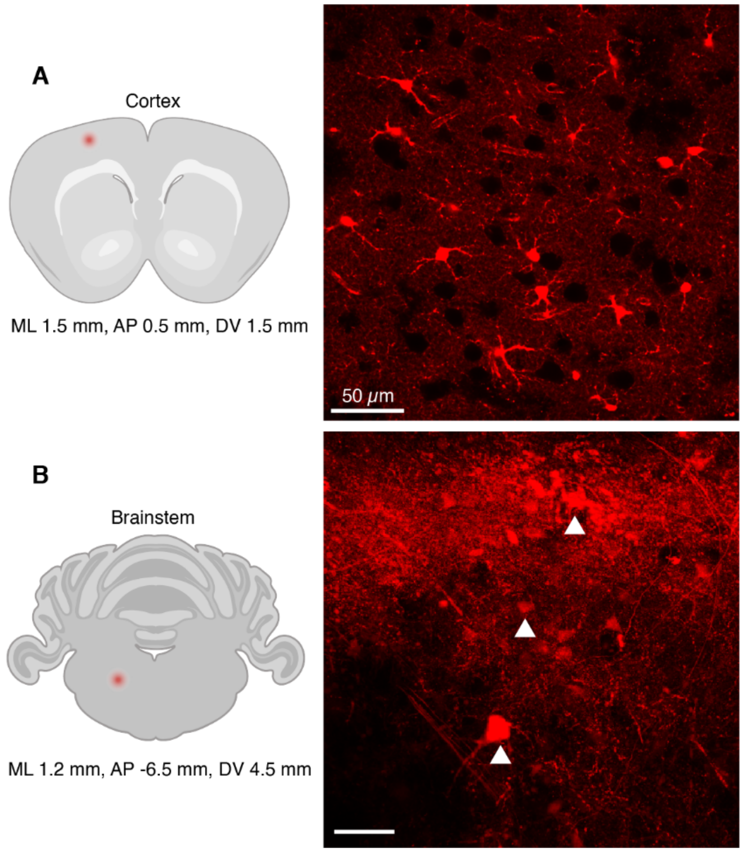


**Fig. S2. Atto 643 labels astrocytes in the cortex but not in the brainstem.** Atto 643 (50 μM) was independently injected into the cortex (**A**) and brainstem (**B**), with stereotaxic coordinates illustrated in the left panel. Acute brain slices were then prepared for two-photon imaging. Right panels: representative images show robust astrocyte labeling by Atto 643 in the cortex; In contrast, no astrocytic labeling was observed in the brainstem, where the dye either accumulated extracellularly or nonspecifically labeled cells with neuronal morphology (white arrowheads in **B**).
